# *Listeria monocytogenes* InlP interacts with afadin and facilitates basement membrane crossing

**DOI:** 10.1101/242222

**Authors:** Cristina Faralla, Effie E. Bastounis, Fabian E. Ortega, Samuel H. Light, Gabrielle Rizzuto, Salvatorre Nocadello, Wayne F. Anderson, Jennifer R. Robbins, Julie A. Theriot, Anna I. Bakardjiev

**Author notes:** These authors contributed equally (Cristina Faralla, Effie E. Bastounis). Current address: VIR Biotechnology, San Francisco, California. Current address: AbCellera Biologics Inc., Vancouver, BC, Canada.

## Abstract

During pregnancy, the placenta protects the fetus against the maternal immune response, as well as bacterial and viral pathogens. Bacterial pathogens that have evolved specific mechanisms of breaching this barrier, such as *Listeria monocytogenes*, present a unique opportunity for learning how the placenta carries out its protective function. We previously identified the *L. monocytogenes* protein Internalin P (InlP) as a secreted virulence factor critical for placental infection (1). Here, we show that InlP, but not the highly similar *L. monocytogenes* internalin Lmo2027, binds to human afadin (encoded by *AF-6*), a protein associated with cell-cell junctions. A crystal structure of InlP reveals several unique features, including an extended leucine-rich repeat (LRR) domain with a distinctive Ca2+-binding site. Despite afadin’s involvement in the formation of cell-cell junctions, MDCK epithelial cells expressing InlP displayed a decrease in the magnitude of the traction stresses they could exert on deformable substrates, similar to the decrease in traction exhibited by *AF-6* knock-out MDCK cells. *L. monocytogenes ΔinlP* mutants were deficient in their ability to form actin-rich protrusions from the basal face of polarized epithelial monolayers, a necessary step in the crossing of such monolayers (transcytosis). A similar phenotype was observed for bacteria expressing an internal in-frame deletion in *inlP* (*inlP* DLRR5) that specifically disrupts its interaction with afadin. However, afadin deletion in the host cells did not rescue the transcytosis defect. We conclude that secreted InlP targets cytosolic afadin to specifically promote *L. monocytogenes* transcytosis across the basal face of epithelial monolayers, which may contribute to the crossing of the basement membrane during placental infection.

## INTRODUCTION

During pregnancy, the consequences of placental infection can be severe, ranging from maternal sepsis to miscarriage, and can lead to pre-term birth and lifelong disability (2). Fortunately, such infections are relatively rare – which stands as a testament to the strength of the feto-maternal barrier. Despite serving such an important function, the molecular, cellular and histological components of feto-maternal barrier have only just begun to be elucidated. Because the barrier is so effective at preventing infection, pathogens that do manage to cross it must have evolved strategies of countering host defenses and thus provide a unique opportunity for addressing the mechanistic features that make the feto-maternal barrier so formidable (3).

*Listeria monocytogenes* is a well-characterized food-borne pathogen capable of placental crossing and is thus ideal for probing this barrier (4). In the healthy, non-pregnant adult, it causes gastrointestinal illness, but in immunocompromised individuals meningitis can result, and in pregnant women, sepsis, spontaneous abortion, preterm labor, infant brain damage and death are possible outcomes (5). After ingestion, *L. monocytogenes* infects epithelial cells of the intestine, mainly via interactions between the bacterial surface protein Internalin (encoded by the gene *inlA*) and the host cell receptor E-cadherin, which is exposed at the tips of intestinal villi (6-8). After uptake by an intestinal epithelial cell, the bacterium escapes the resulting phagosome and replicates in the host cytoplasm, where it induces actin polymerization, forming actin comet tails that drive its rapid motility (9). Upon reaching the host plasma membrane, *L. monocytogenes* can form a membrane-bound protrusion that pushes its way into a neighboring cell. Engulfment of the protrusion by the neighboring cell leaves the bacterium in a double-membrane vacuole from which it escapes, resetting its life cycle (9, 10). From the initial site of infection in the intestinal epithelium, *L. monocytogenes* can spread via actin-based motility into immune cells, which facilitate spread throughout the host while protecting the pathogen from humoral immune defenses (11, 12). Actin-based cell-to-cell spread is thought to contribute to the ability of *L. monocytogenes* to evade the host immune system and to penetrate a variety of protective barriers in the host (13), including the blood-brain barrier and the feto-maternal barrier in the placenta (14-16).

A clue to the protective nature of the feto-maternal barrier lies in the fact that almost all pathogens that cross it are known to have an intracellular aspect to their life cycle—they move *through* cells from mother to fetus (3). Two interfaces are available; each is unlike any other part of the mammalian body. The first, where fetal extravillous trophoblasts (EVTs) anchor the placenta in uterine decidua, has a unique immune environment (17). The trophoblasts possess innate immune defense properties shown to prevent bacterial infection (18) and restrict growth of intracellular *L. monocytogenes* (19). The second, much more extensive interface is composed of fetal syncytiotrophoblasts (STB) which form a vast, thin multinucleate layer without cell-cell junctions, bathed in maternal blood. This is the site of gas and nutrient/waste exchange. Underlying the STB is a second, single-celled layer of individual, self-renewing cytotrophoblasts (CTBs) that periodically fuse with the STB to allow its growth. Our previous work has shown that the STB’s lack of cell-cell junctions (20) and the syncytial stiffness generated by dense networks of actin filaments (21) act as significant deterrents to STB infection by pathogens. Rather, infection of human placental organ cultures suggests that the first interface, where EVTs contact uterine decidua, is the preferred route of placental infection by *L. monocytogenes*, which then spreads via actin-dependent cell-to-cell spread to the CTB monolayer underlying the STB (20). But even once *L. monocytogenes* has reached the subsyncytial CTBs, most bacterial movement occurs laterally from cell to cell within the monolayer, and it only rarely transcytoses in a direction perpendicular to the monolayer to colonize the fetal stroma beneath it (20).

Recently, we used an unbiased genetic screen in a pregnant guinea pig model of listeriosis to identify *L. monocytogenes* genes that contribute specifically to infection of the placenta. (1). One gene identified in this screen encodes InlP, a member of the internalin family of proteins. Pathogenic *L. monocytogenes* contain genes for 25 known members of this protein family, which share a common overall structure that includes a secretory signal sequence at the N terminus and a large leucine-rich repeat domain, a motif frequently implicated in protein-protein interactions (22). The two best-characterized internalins, InlA and InlB, contribute to *L. monocytogenes* invasion of epithelial cells and hepatocytes by binding to their cognate host cell surface receptors, E-cadherin and c-Met respectively (6, 23). While most internalins are predicted to be anchored to the bacterial cell surface via attachment to the peptidoglycan cell wall or to lipoteichoic acids, four of them, including InlP, lack obvious anchoring domains and are predicted to be secreted by the bacterium (24). In order to address the mechanism by which InlP assists *L. monocytogenes* in crossing the feto-maternal barrier, we set out to find host protein binding partners. In this work, we identify the host cell cytoplasmic protein afadin as a major binding partner for InlP, and demonstrate that the InlP-afadin interaction specifically enhances *L. monocytogenes* transcytosis—that is, the ability of *L. monocytogenes* to exit from the basal face of an infected polarized epithelial monolayer, consistent with its potential contribution to placental infection.

## MATERIALS AND METHODS

### Study approval

<u>Human subjects:</u> This study was conducted according to the principles expressed in the Declaration of Helsinki. The study was approved by the Institutional Review Board at the University of California, San Francisco, where all experiments were performed (CHR# 11-05530). All patients provided written informed consent for the collection of samples and subsequent analysis.

### Recombinant InlP expression and purification

All chemicals were purchased from Sigma-Aldrich unless otherwise stated. Center for the Structural Genomics of Infectious Diseases (CSGID) standard protocols were used for cloning, over-expression, and purification of InlP (25, 26). Briefly, InlP (from residue 31-388) and Lmo2027 (from residue 31-367) were cloned into the pMCSG7 expression vector (http://bioinformatics.anl.gov/mcsg/technologies/vectors.html).

Following transformation into the BL21 (DE3) Magic *E. coli* strain, cells were grown in M9-selenomethionine medium or Terrific Broth (TB) at 37 °C up to an OD_600_ of 1. At that point, the temperature was reduced to 25 °C and protein over-expression was induced by the addition of isopropyl-1-thio-D-galactopyranoside (IPTG) to a final concentration of 1 mM. After 16 h, cells were harvested by centrifugation, suspended in a buffer containing 10 mM Tris-HCl pH 8.3, 500 mM NaCl, 10% glycerol, and 1 mM Tris (2-carboxyethyl) phosphine hydrochloride (TCEP) and lysed by sonication. InlP was purified by Ni-NTA affinity chromatography and eluted with a buffer containing 10 mM Tris-HCl pH 8.3, 500 mM NaCl, 1 mM TCEP. The His-tag was cleaved overnight at 4°C incubating the enzymes with His-tagged TEV protease. The protein samples were then reloaded onto the nickel column and the flow-through was collected. At this point, the protein was concentrated using Vivaspin centrifugal concentrators (GE Healthcare Life Sciences) and both its size and purity were checked by SDS-PAGE.

### InlP and Lmo2027 structure determination

Sitting drop crystallization plates were set up at room temperature. InlP crystals were obtained with a 1:1 mixture of 8.5 mg/mL InlP in 10 mM Tris pH 8.3, 1 mM TCEP and condition D11 from the Qiagen PACT crystallization screen (0.2 M Calcium chloride, 0.1 M Tris pH 8, 20% (w/v) PEG 6000). Lmo2027 crystals were obtained with a 1:1 mixture of 12.6 mg/mL Lmo2027 and 10 mM Tris pH 8.3, 0.5 M NaCl, and 1 mM TCEP and condition D4 from the Qiagen Classics II crystallization screen (0.1 M Citric acid (pH 3.5) and 20% (w/v) PEG 3350). Harvested crystals were transferred to the mother liquor before being frozen in liquid nitrogen. Diffraction data were collected at 100o K at the Life Sciences Collaborative Access Team at the Advance Photon Source, Argonne, Illinois (APS BEAMLINE 21-ID-G). All structural work was performed by the CSGID (25). Data were processed using HKL-2000 for indexing, integration, and scaling (27). A selenomethionine derivative of InlP was used to phase the structure by single-wavelength anomalous diffraction. Lmo2027 was phased by molecular replacement, using the InlP structure as a starting model. Structure was refined with Refmac (28). Models were displayed in Coot and manually corrected based on electron density maps (29). All structure figures were prepared using PyMOL Molecular Graphics System, Version 1.3 (Schrödinger, LLC).

### Isothermal titration calorimetry (ITC)

For ITC experiments, InlP was prepared by overnight dialysis in ITC buffer (50 mM HEPES pH 7.5 and 150 mM NaCl). Experiments were performed on the MicroCal ITC200 instrument (GE Healthcare). InlP was loaded into the cell at 0.12 mM and 25° C. Calcium chloride titrant (3.13 mM) dissolved in leftover dialysis buffer was loaded into the syringe. Syringe rotation was set at 1000 RPM, with titrant injections spaced at 2 min intervals. The initial 0.2 μL injection was excluded from the data and the nineteen subsequent 2.0 μL injections were used for data analysis. Binding parameters were obtained by fitting isotherms using the Origin 7 (OriginLab, Northampton, MA) software package.

### Yeast two-hybrid, mass spectrometry analysis and pull-down assay

Yeast two-hybrid analysis was performed by Hybrigenics (France). Briefly, InlP (aa 31-388) was used to screen a human placenta cDNA library. Clones expressing proteins with positive interactions with InlP were isolated and sequenced.

For the GST-pull down experiments bait proteins were prepared as follows: Cloning: *inlP* gene (from residue 31-388, oligo Fw2470 EcoRI-pGex CCGAATTCC GCTTCTGATTTATATCCACTACCT, Rv2470 XhoI-pGex aGCTCGAG TCATTAATAGTTACATTCCAATCATAAGAG) or Lmo2027 (from residue 31-367, oligo Fw2027 EcoRI-pGex CCCCGAATTCCGCATCCGATTTATATCCACT ACC, Rv2027 XhoI-pGex GAAGCTCGAGCTACTAATTAACCGTAAGTACCC) were cloned in pGEX-4T vector (GE Healthcare) using restriction enzymes following transformation in BL21 DE3 *E. coli* strain. InlPΔLRR5, InlPΔLRR7 and InlPΔLRR8, lacking the LRR5 (Δamino acids 174-195), the LRR7 motifs (Δamino acids 218-239) and the LRR8 (Δamino acids 240-261) respectively were generated using Site Directed Mutagenesis (SDM) on the *inlP*-pGEX-4T vector following In- Fusion^®^ HD Cloning Kit protocol (Takara Bio USA, Inc.) using the following primers: ΔLRR5-fw:GATTTCACTGGAATGCCTATTCCTTTACTTATTACGTTGGATCTAAG, ΔLRR5-rv: CGTAATAAGTAAAGGAATAGG CATTCCAGTGAAATCAGGGATAGA; ΔLRR7-fw CCTGATTTTCAAAATTTACCT AAATTAACTGATTTAAATTTAAGAC, ΔLRR7-rv: AGTTAATTTAGGTAAATTTTGAAAATCAGGAATAGTTGTCA, ΔLRR8-fw: CCTGATTTTCAAA ATAACTTACCTAGTTTAGAATCCTTAAACT ΔLRR8-rv: TAAACTAGGTAAGTTATTTTGAAAATCAGGTGTATTGGTTAA. Baits purification: BL21 DE3 *E. coli* carrying pGEX-4T, pGEX-4T-*2027*, pGEX-4T-*inlP*, pGEX-4T-*inlP*Δ*LRR5*, pGEX-4T-*inlP*Δ*LRR7* or pGEX-4T-*inlP*Δ*LRR8*, were grown until OD 0.6 in 250 mL of LB supplemented with 100 µg/mL ampicillin at 37 °C and 180 rpm. GST or GST-InlP/2027/ΔLRR5/ΔLLR7/ΔLRR8 expression were then induced with 0.2 mM IPTG for 4 h at 30 °C 180 rpm. Pellet from 250 mL of culture was lysed with 35 mL of cell lytic express (Sigma) and the soluble fraction was incubated overnight with 5 mL of Glutathione Sepharose™ 4B (GE Healthcare) under constant agitation. GST-Glutathione Sepharose™ 4B or GST- InlP/2027/ΔLRR5/ΔLLR7/ΔLRR8/ - Glutathione Sepharose™ 4B were then collected and washed with 100 mL of PBS at 500 x g and stored at 4 °C in Tris-HCl 50 mM, NaCl 50 mM pH 8.

For preparation of protein extracts for pull-down experiments with human placenta: 2.3 grams of placental villi (~21 weeks gestational age) were lysed in 10 mL of M-PER™ (Thermo Fisher Scientific) supplemented with one tablet of complete Protease Inhibitor Cocktail (Roche) and PhosSTOP phosphatase inhibitor (Roche). The sample was homogenized for 30 sec, sonicated for 4 min and incubated for 1 h on ice. The soluble fraction was collected after centrifugation 14000 x g for 20 min at 4 °C, split in two and incubated overnight under constant agitation with 1 mL of GST-Glutathione Sepharose™ 4B, GST-InlP- Glutathione Sepharose™ 4B or GST-2027- Glutathione Sepharose™ 4B respectively (prepared previously). The next day, washes were performed with 50 mL PBS and elution fractions were collected with 3 mL Tris-HCl 50 mM, NaCl 50 mM free glutathione 10 mM, pH 8. Elution fractions were run on 10% gel (NuPAGE™ Novex™ 10% Bis-Tris Protein Gels) and analyzed by Western blot, Coomassie blue staining (SimplyBlue™ SafeStain, Life Technologies) or sent to Taplin Mass Spectrometry Facility at Harvard Medical school for protein identification from SDS-PAGE gel (Supplementary Table 2).

In preparation of protein extracts for pull-down experiments with MDCK cells: one 75 mm flask of MDCK or MDCK *AF-6*^-/-^ cells polarized for three days was lysed in 180 µL of cell lytic M supplemented with complete Protease Inhibitor Cocktail (Roche). The soluble fraction was split in two and incubated overnight under constant agitation with 20 µL of GST-Glutathione Sepharose™ 4B or GST-InlP/2027/ΔLRR5/ΔLRR7/- Glutathione Sepharose™ 4B respectively (prepared previously). The day after beads were washed with 2 mL of Tris-HCl 50 mM, NaCl 50 mM, free glutathione 10 mM, pH 8.0. The bound proteins were eluted by boiling the beads in SDS sample buffer (NuPAGE^®^ sample reducing agent 10X and NuPAGE^®^ LDS sample buffer 4X) for 30 min and analyzed by Western Blot or Coomassie blue staining (SimplyBlue™ SafeStain, Life Technologies) or sent to Taplin Mass Spectrometry Facility at Harvard Medical school for protein identification (Supplementary Table 2 and Fig. 3D). Raw mass spectrometry data were then further analyzed to determine differences in binding of proteins to InlP-GST versus InlPΔLRR5- GST. First proteins identified by only 1 peptide in 1 or 2 samples were excluded from further analysis. Then proteins found to bind to GST alone were also excluded from further analysis. Finally proteins that showed 0 intensity for either InlP-GST or InlPΔLRR5-GST were also excluded from the analysis. The resultant data and analysis are depicted in Fig. 3D and Fig. S4.

To detect afadin by Western Blot analysis, elution fractions were analyzed on 4-12% gels (NuPAGE™ Novex™ 4-12% Bis-Tris Protein Gels) run on ice in MOPS buffer (NuPAGE^®^ MOPS SDS Running Buffer) for 3 h at 200 V and transfer was performed at 100V for 2 h. Primary staining using mouse anti-afadin antibody (BD Transduction Laboratories™, 1:2000) and mouse anti-β-actin antibody (SIGMA, 1:1000) were incubated overnight at 4 °C. Secondary antibody goat anti-mouse IgG-HRP (1:10000, Santa Cruz Biotechnology’s) was incubated for 1 h at room temperature. Protein detection was performed using ECL Western Blotting Substrate™ (Pierce).

### Human tissue collection and immunofluorescence

For human placental organ cultures, placentas from elective terminations of pregnancy (gestational age 4 to 8 weeks) were collected and prepared as previously described (30). Briefly, fragments from the surface of the placenta were dissected into 3-6 mm tree-like villi, placed on Matrigel (BD Biosciences)-coated Transwell filters (Millipore, 30-mm diameter, 0.4 μm pore size) and cultured in Dulbecco’s modified Eagle-F12 medium (DMEM-F12; 1:1, vol/vol) supplemented with 20% FBS, 1% L-glutamine and 1% penicillin/streptomycin (Invitrogen). Organ cultures were fixed for 15-30 min in 4% paraformaldehyde at room temperature, flash frozen in Tissue-Tek O.C.T. Compound (Sakura Finetek, Torrance, CA). Sections were cut at 7 μM, permeabilized for 5 min in ice-cold acetone, dried, rehydrated in PBS, and blocked with Background Buster (Innovex Biosciences, Richmond, CA). Primary staining with mouse anti-afadin antibody (BD Transduction Laboratories™, 1:2000) was incubated for 2 h at room temperature in PBS with 1% BSA and 5% normal goat serum (Jackson ImmunoResearch Laboratories). Secondary staining with Alexa-Fluor® 594 goat anti-mouse (1:500, ThermoFisher Scientific) was incubated for 1 h at room temperature, nuclei stained with DAPI (Affymetrix), and sections mounted in Vectashield Mounting Medium (Vector Laboratories). Stitched high-power images were acquired on a Nikon Ti-E epifluorescent microscope with DS-Ri2 camera and NIS-Elements 4.30 software (Nikon Corporation).

### Mammalian cell line tissue culture

MDCK (Madin Darby Canine kidney cell line, ATCC), MDCK-AF6-/- (a generous gift from Denise Marciano, UT Southwestern) cells were grown in Eagle’s MEM (Minimal Essential Medium) with Earle’s Balanced Salt Solution (BSS) supplemented with 10% FBS. Invasion assays and growth curves were performed as previously described (31). MDCK cells expressing a humanized version of InlP were generated as follows: the IRES-GFP sequence was amplified from the pIRES-EGFP-puro plasmid (Addgene) using oligo fw-IRESGFP-EcoRI: GGAATTC<u>GAATTC</u>TAAGCCCCTCTCCCTCCCCCC and rv-IRESGFP-BamHI GGAATTC<u>GGATCC</u>TTACTTGTACAGCTC GTCCATGCCG and cloned into the plasmid PCR2.1. The *inlP* gene was codon optimized for expression in mammalian cells (h*inlP*) and the NheI-h*inlP*-Flag-EcoRI construct was synthesized by GeneWiz. Using the restriction enzymes NheI and EcoRI the NheI-h*inlP*-Flag-EcoRI construct was then cloned into PCR2.1-IRES-GFP. The construct NheI-h*inlP*-Flag-EcoRI-IRES-GFP-BamHI was then cut from PCR2.1 using the restriction enzymes NheI-BamHI and cloned in pCW-Cas9-Blast (Addgene) plasmid by replacing the Cas9 gene and generating the plasmid pCW- h*inlP*-Flag-IRES-GFP. UCSF ViraCore (http://viracore.ucsf.edu/) was used to generate a Lentivirus strain carrying pCW- h*inlP*-Flag-IRES-GFP, which was used to transduce MDCK or MDCK *AF-6*-/- cells. Viral transduction was performed in 12-well plates as follows: freshly plated MDCK cells were allowed to adhere for 1 h at 37 °C (4x10^4^ cells/well), next nearly all media was removed and 350 μl of viral supernatant was added to each well, followed by overnight incubation at 37 °C. The next morning 1 mL fresh media was added to each well without removing the viral supernatant and cells were grown for 48-72 h, followed by puromycin selection (5 μg/mL) for another 4 days. Protein expression was analyzed by Western Blot +/- addition of doxycycline 1 μg/mL. Primary staining: mouse anti-flag antibody (1:5000), and mouse anti-β-actin antibody (SIGMA, 1:1000), secondary antibody: goat anti-mouse IgG-HRP (1:10000, Santa Cruz Biotechnology). Protein detection was performed using ECL Western Blotting Substrate™ (Pierce).

### Traction force microscopy (TFM)

Single cell TFM assays were performed as previously described (32, 33). Two layered polyacrylamide hydrogels, the upper of which contained 0.04% carboxylate-modified red latex beads of 0.1 μm in diameter (FluoSpheres; Molecular Probes) were prepared and activated via SulfoSanpah crosslinker (Thermo Fischer Scientific, 22589) as previously described (32, 33). Hydrogels were attached to 24-well glass bottom plates (MatTek) to enable monitoring of multiple conditions simultaneously. The stiffness of the hydrogels was 5 kPa, achieved with a 5% final concentration of acrylamide (Sigma, A4058) and 0.15% final concentration of bis-acrylamide (Fisher, BP1404-250). Activated hydrogels were coated with collagen I (Sigma-Aldrich, C3867) overnight at 4 °C. Hydrogels were washed with PBS the next day and equilibrated with cell media for 30 min at 37 °C prior to addition of cells.

For single cell TFM experiments, 2 mL containing 10x10^5^ cells were added on six-well TC polystyrene plates for 24 h. Upon seeding 1 μg/mL doxycycline was added to the wells where InlP expression was desired. The next day, cells were lifted via 0.25% trypsin-EDTA, washed once in PBS and then cell pellets were re-suspended in media and 0.5x104 cells in 1 mL were added per hydrogel to achieve single cell attachment. Cells were allowed to adhere for 3 h at 37 °C before imaging. To measure cell-ECM traction forces in confluent monolayers cells were added to a concentration of 4x10^5^ cells per well directly onto the hydrogels 24 h prior to imaging. Upon seeding 1 μg/mL doxycycline was added to the wells where InlP expression was desired.

Multi-channel time-lapse sequences of fluorescence (to image the beads within the upper portion of the hydrogels) and phase contrast images (to image the cells) were acquired using an inverted Nikon Diaphot 200 with a CCD camera (Andor Technologies) using a 40X Plan Fluor NA 0.60 objective and the MicroManager software package (Open Imaging). In addition, at the beginning and end of the recordings an image of the cells’ nuclei stained with 1 (μg/mL Hoechst and an image of InlP fluorescence were acquired to ensure similar cell densities and similar InlP expression across conditions, respectively. The microscope was surrounded by a cage incubator (Haison) maintained at 37 °C and 5% CO_2_. Images were acquired every 5 min for approximately 4 h. Subsequently, at each time interval we measured the 2D deformation of the substrate at each point using an image correlation technique similar to particle image velocimetry (34). We calculated the local deformation vector by performing image correlation between each image and an undeformed reference image which we acquired by adding 10% SDS at the end of each recording to detach the cells from the hydrogels. We used interrogation windows of 32 x 16 pixels (window size x window overlap) for cells forming a monolayer or 32 x 8 pixels for single cells. For single cell experiments, we used a custom algorithm using MATLAB (MathWorks) to identify the contour of the cells from the phase contrast images (35). We calculated the two-dimensional traction stresses that single cells exert to the hydrogel as described elsewhere (35). We calculated the strain energy (*U_s_*) as the mechanical work imparted by the cell to deform its hydrogel:

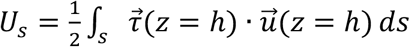, where 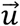 is the measured displacement vector field on the free surface of the hydrogel and ∫*_s_* () *ds* represents a surface integral. For cell monolayer experiments, traction stresses were measured as described above and elsewhere (33, 36).

### *In vitro* transcytosis and invasion assays

To quantify *L. monocytogenes* transcytosis, MDCK cells were seeded on Transwell™ inserts with a 3-*µ*m pore size loaded onto tissue culturetreated 12-well plates (Corning, 3402) at a density of 5x10^5^ cells per insert three days prior to infection. The day before infection, *L. monocytogenes* cultures were inoculated in 3 mL of Brain Heart Infusion (BHI) directly from glycerol stocks and grown at 30 °C for 15 to 16 h without agitation. The day of the experiment, the optical density of the overnight cultures was measured to ensure that the MDCK cells were infected with a comparable multiplicity of infection (MOI) across bacterial strains. For each strain, 1.2 mL was centrifuged at 2000 x g for 5 min, and washed with 1 mL of DPBS. Bacterial pellets were resuspended in 16 mL of MEM. Host cells were washed with MEM once, infected with 0.5 mL of bacterial mix per Transwell, and incubated at 37 °C for 10 min. The unused bacterial mix was serially diluted and plated onto BHI agar plates containing 200 *µ*g/mL streptomycin for bacterial enumeration. Transwells were washed three times with MEM alone, then basal and apical media were both replaced with MEM + 10% FBS, and plates were incubated at 37 °C for 1 h to allow bacterial invasion, followed by replacement with MEM + 10% FBS + 50 *µ*g/mL gentamicin to kill extracellular bacteria, and incubated at 37 °C for 15 min more. To assess subsequent transcytosis, Transwells were washed three times with MEM + 10% FBS, and were then transferred to a new 12-well plate containing 1 mL of fresh MEM + 10% FBS per well. This plate was incubated at 37 °C for 1 h, at which point basal media was collected, vortex vigorously and 100 *µ*L of each sample plated onto BHI agar plates containing 200 *µ*g/mL streptomycin. This step served as a negative control to exclude for damaged monolayers. Meanwhile, the Transwells were transferred to another new 12-well plate containing fresh MEM + 10% FBS per well. After an additional 3 h at 37 °C, basal media was plated on BHI agar plates containing 200 *µ*g/mL streptomycin as before, this time to assess true transcytosis.

To quantify *L. monocytogenes* invasion, MDCK ce3^rd^ were seeded onto tissue culture-treated 12-well plates at a density of 5x10^5^ cells per well three days prior to infection. MDCK cells were then infected with an *ΔactA L. monocytogenes* strain expressing mTagRFP under the actA promoter (37), which becomes transcriptionally active upon entry into the host cell cytoplasm. Invasion was assayed with flow cytometry as previously described (38).

## RESULTS

### InlP binds afadin

We attempted to find host binding partners for InlP in placental cells using a yeast two-hybrid system to screen a human placenta library (39) (Supplementary Table 1). In parallel, we used InlP fused to glutathione-S-transferase (InlP-GST) and a mass spectrometric approach to pull-down and identify InlP binding partners in human placental tissue extracts (Supplementary Fig. 1 and Supplementary Table 2). Both methods should in principle be able to detect both host cell surface and cytoplasmic binding partners for InlP. While each of these approaches identified multiple candidates, the protein afadin, encoded by the *MLLT4/AF-6* gene, was the only one identified by both. Afadin is an F-actin-binding protein that also binds to the cytoplasmic domain of the cell-cell adhesion molecule nectin, as well as to a wide variety of signaling proteins including Src and multiple members of the Ras superfamily (40). Afadin has been shown to contribute to the formation of cell-cell junctions during mouse embryonic development (41, 42) and to apical constriction of adherens junctions during *Drosophila* morphogenesis (43). In addition, afadin has been proposed to play multiple roles in cell polarization, migration, differentiation, and oncogenesis (40).

We confirmed specific binding of InlP to afadin by pull-down experiments followed by Western blot analyses (Fig. 1A and 1B). Elution fractions obtained after pull-down using an InlP- GST fusion protein incubated with cell lysates from MDCK, MDCK *AF-6*^-/-^ or human placental villi were analyzed. Using anti-afadin antibodies, we detected afadin signal in the input fraction and in the elution fractions after incubation of InlP-GST with protein extract from MDCK cell lines (Fig. 1A) or from human placental villi (Fig. 1B). No afadin was detected in the elution fractions when the GST control was used as bait (Fig. 1A and 1B). Confirming the specificity of our antibodies, Western blot analysis on pull-down experiments performed using protein extract from the afadin knock-out MDCK cell line did not show a detectable signal (Fig. 1A).

**Figure 1.**
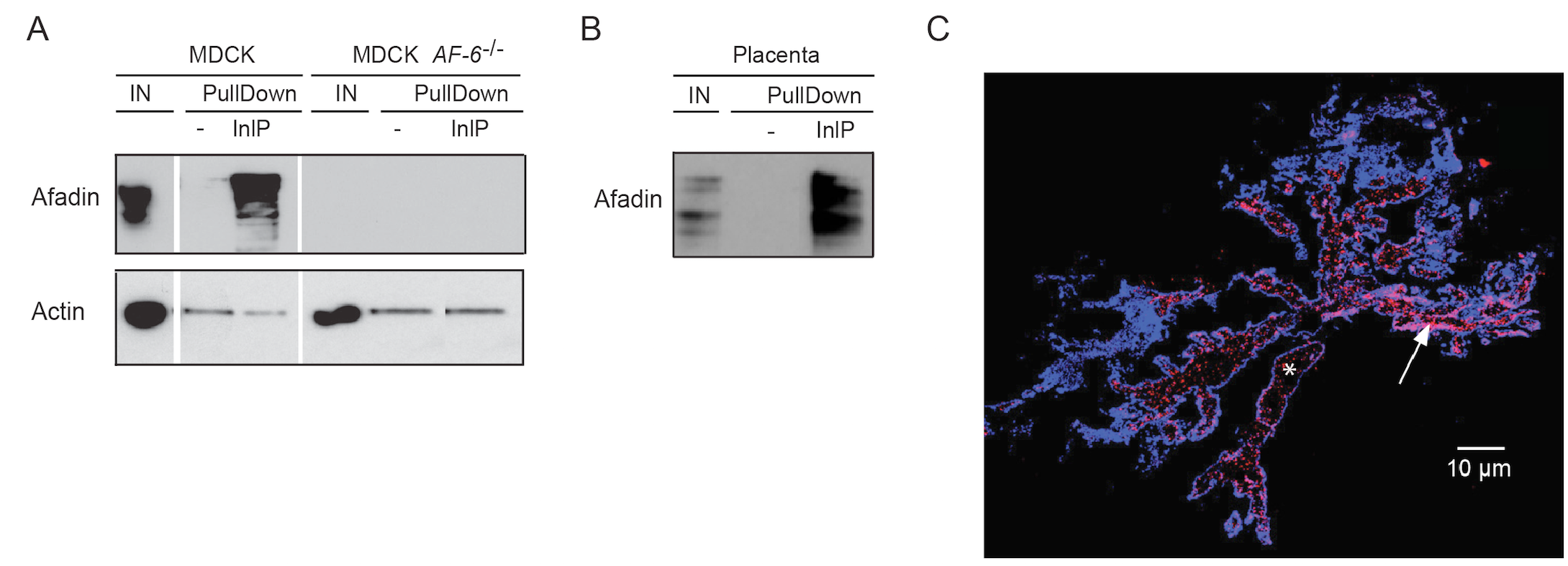
Bacterial effector InlP binds host adaptor protein afadin. (A-B) InlP-GST fusion protein (InlP) or GST protein alone (-) bound to glutathione-Sepharose resin were incubated overnight with protein extracts from (A) MDCK or MDCK *AF-6*^-/-^ cells or (B) human placenta. Input (IN) and elution fractions (EF) were analyzed by Western blot with anti-afadin antibodies. Actin: loading control. (C) Immunofluorescence of human placental villi stained for afadin (red) and DAPI (blue), white bar =10 *µ*m (right). Asterisk labels a portion of the stroma, and the arrow points to some of the CTBs.

To determine the amount and spatial distribution of afadin present in human placental villi, we performed immunofluorescence microscopy staining for afadin and observed that this protein is very abundant in the placenta (Fig. 1C). Localization was primarily observed at the CTB layer just beneath the syncytium, with small amounts in the stroma (see asterisk and arrow in Fig. 1C). The high abundance of this potential InlP binding partner in the placenta, combined with our previous report that InlP plays an important role in *L. monocytogenes* infection at the maternal-fetal interface (1), are consistent with the hypothesis that InlP-afadin interactions are important for the pathogenesis of placental infections.

### Lmo2027 is a structural paralog of InlP that does not bind afadin

The internalins that are predicted to be secreted by *L. monocytogenes* include InlC (Lmo1786), InlP (Lmo2470), Lmo2027 and Lmo2445 (24). Of these, only InlC has been extensively studied, but its primary sequence is not particularly similar to that of InlP (24% sequence identity). The two other internalins predicted to be secreted, Lmo2027 and Lmo2445, have not yet been characterized. Pairwise analyses of amino acid sequences indicated that InlP and Lmo2445 are only 25% identical and 40% similar; however, InlP and Lmo2027 are much less divergent, with 65% identity and 77% similarity (Fig 2A). Moreover, InlP and Lmo2027 share a similar C-terminal domain that is not found in any of the other internalin family members (24).

**Figure 2.**
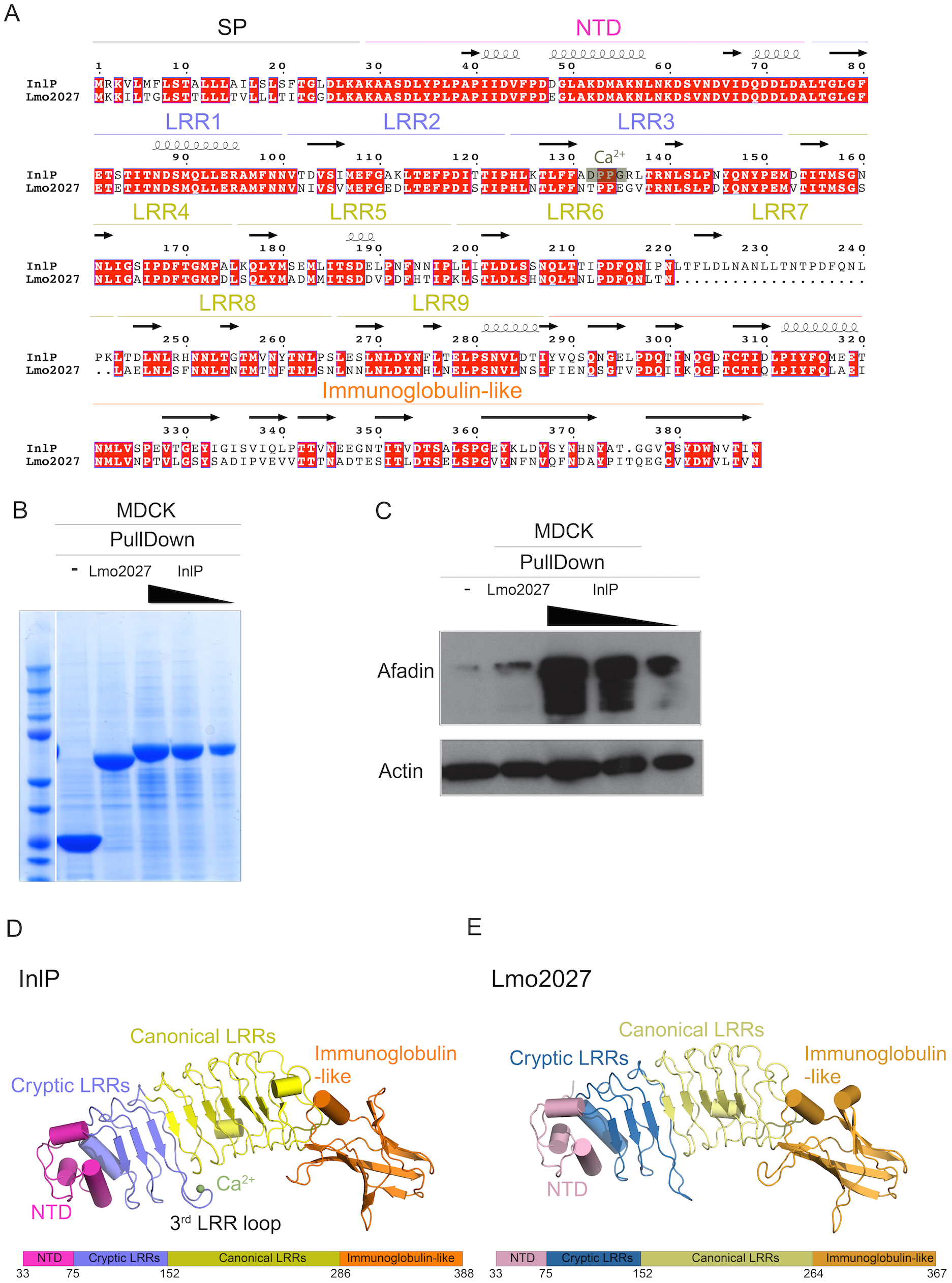
Distinct features of InlP as compared to Lmo2027. (A) Pairwise sequence alignment of InlP and Lmo2027 with conserved amino acids highlighted in red. The LRRs and calcium binding site on InlP are noted. Arrows signify ß-sheets and coils correspond to α-helices. (B) Loading control of pull down experiments on Lmo2027 by Coomassie blue staining. GST protein alone (-), Lmo2027-GST fusion protein (2027) or InlP-GST fusion protein (InlP) bound to glutathione-Sepharose resin used as bait for pull-down experiments with protein extracts from MDCK cells. The elution fraction from pull-down samples using InlP-GST (InlP) as bait were sequentially diluted 5-fold before SDS analysis; undiluted elution fractions were analyzed from pull-down samples using GST (-) or Lmo2027 (2027). Data shown are Coomassie staining and the most abundant band in each lane represents the bait. (C) GST fusion proteins or GST protein alone (-) bound to glutathione-Sepharose resin were incubated overnight with protein extracts from MDCK cells, and elution fractions were analyzed by Western blot with anti-afadin antibodies. The following GST fusion proteins were used: GST-Lmo2027 (Lmo2027) and GST- InlP (InlP). Undiluted elution fractions were analyzed with the exception of the elution fractions marked by black triangles: elution fractions from the experiments using GST-InlP as bait were sequentially diluted by 5-fold before western blot analysis. Actin: loading control. (D) Crystal structure of InlP with schematic depiction of domain layout below. (E) Crystal structure of Lmo2027 with color-coded domains, like in (D).

Because of its high degree of sequence similarity to InlP, we decided to test whether Lmo2027 also binds afadin. As before, we performed pull-down assays with MDCK cell lysates, followed by Western blots on elution fractions using Lmo2027-GST or InlP-GST fusion proteins as bait (Fig. 2B, C). Unlike InlP-GST, Lmo2027-GST did not enrich the afadin signal in the elution fractions significantly more than the GST control. Even when diluted 10-fold, InlP-GST still pulled down considerable afadin in comparison to the GST control and Lmo2027-GST. These results show that, despite the relatively high sequence similarity of the two proteins, InlP, but not Lmo2027, binds afadin.

To identify structural distinctions that might account for differences in afadin binding affinity, we obtained 1.4 Å and 2.3 Å crystal structures of InlP and Lmo2027, respectively (Supplementary Table 3). Analysis of the structures reveals that InlP and Lmo2027 generally resemble previously characterized internalins (Fig 2D, E). Both proteins possess an N-terminal domain that adopts the helical structure characteristic of internalins. This is followed by a concave leucine rich-repeat (LRR) domain made up of 9 LRRs in InlP and 8 LRRs in Lmo2027 that can be further subdivided into a “cryptic” portion consisting of 3 LRRs, which were not anticipated to adopt an LRR-fold based upon bioinformatics analyses, and a “canonical” portion consisting of 6 LRRs in InlP and 5 LRRs in Lmo2027. Finally, the C-terminal domain in both proteins assumes an immunoglobulin-like fold.

In particular, two features distinguish InlP and Lmo2027 from each other and from previously characterized internalins. First, the additional LRR in InlP extends the LRR domain in this protein as compared to Lmo2027 and increases the distance between the N-terminal and Ig-like domains by ∽4 Å. Such a fundamental distinction in the dimensions of the internalin could affect whether a prospective binding partner could dock to and bind the LRR. Second, the concave surface of previously crystallized internalin LRRs present an unbroken β-sheet (24), but in InlP and Lmo2027 the third cryptic LRR (3^rd^ LRR loop) protrudes, adopting a twisted conformation that disrupts the continuity of this surface. Since this region in other LRR proteins is generally involved in engaging the protein binding partner (22, 44), the distinct 3^rd^ LRR loop seems well situated to play a role in conferring binding specificity.

Unexpectedly, a calcium ion (from the crystallization solution) bound to the center of the 3^rd^ LRR loop structure in InlP, but not Lmo2027 (Fig. 2D, E; Supplementary Fig. 2A), suggesting that a calcium-stabilized loop conformation might be relevant for InlP protein-protein interactions. Two of the residues specifically involved in coordination of the calcium ion in the InlP crystal structure, D132 and G135, are not conserved in Lmo2027 (changed to T and E respectively). To examine the interaction between InlP and calcium, we performed isothermal calorimetry experiments, and found that calcium binds with a relatively low affinity of 35 μM (Supplementary Fig. 2B).

### InlPΔLRR5 has reduced afadin binding affinity

Based on the alignment shown in Fig. 2A, it is evident that the main differences between InlP and Lmo2027 are amino acid differences in that cluster to the C-terminal domain, as well as the absence of 22 amino acid residues of Lmo2027 corresponding to one of the LRR motifs of InlP. Given the high sequence similarity of the 6 “canonical” LRR motifs, it is difficult to definitively identify which one is missing in Lmo2027, even if the pairwise alignment between InlP and Lmo2027 suggests that the missing LRR motif might be LRR7. Therefore, we decided to systematically delete individual LRR motifs in the InlP protein and test the effect of these deletions on the protein’s ability to bind afadin. We successfully generated three InlP deletion mutants—InlPΔLRR5-GST, InlPΔLRR7-GST and InlPΔLRR8-GST—lacking the LRR5 (Δamino acids 174-195), the LRR7 (Δamino acids 218-239) and the LRR8 (Δamino acids 240-261) motifs, respectively. Each of these was then used as bait in pull-down experiments with MDCK cell lysates, followed by Western blot analysis of the elution fractions to detect afadin.

Resins prepared using *E. coli* lysate expressing InlPΔLRR7-GST (Fig. 3A) pulled down afadin from MDCK cell lysates with an efficiency roughly comparable to that observed for InlP- GST (Fig. 3B). In contrast, we detected lower amounts of afadin in the pull-down fraction when lysate from *E. coli* expressing the InlPΔLRR5-GST fusion protein was used to prepare the resin (Fig. 3B). As the expression level of the InlPΔLRR5-GST fusion protein in *E. coli* appeared to be somewhat lower than that of either InlP-GST or GST alone (Fig. 3A), we also performed a Western blot comparing the elution fractions after five-fold dilution for both InlP-GST and GST with no dilution for the InlPΔLRR5-GST elution fraction. Even compared to diluted InlP-GST, substantially less afadin was detected in the sample for InlPΔLRR5-GST (Fig. 3C). This suggests that deletion of the LRR5 motif decreases the affinity of afadin by at least ten-fold. The solubility of the InlPΔLRR5-GST recombinant fusion protein was confirmed by dynamic light scattering analysis (data not shown), consistent with the hypothesis that this reduction in apparent binding affinity is due to specific structural alterations in the LRR domain. The InlPΔLRR8-GST was expressed in *E. coli* lysates at ∽30% of InlP-GST levels prepared under similar conditions (Supplementary Fig. 3A). While we did observe reduced afadin binding by resins prepared using *E. coli* lysates containing this fusion protein as compared to native InlP-GST (Supplementary Fig. 3B), this decrease was approximately proportional to the reduction in expression level. Overall this analysis suggested that InlPΔLRR5 represents a useful mutant for interrogating the InlP-Afadin interaction.

**Figure 3.**
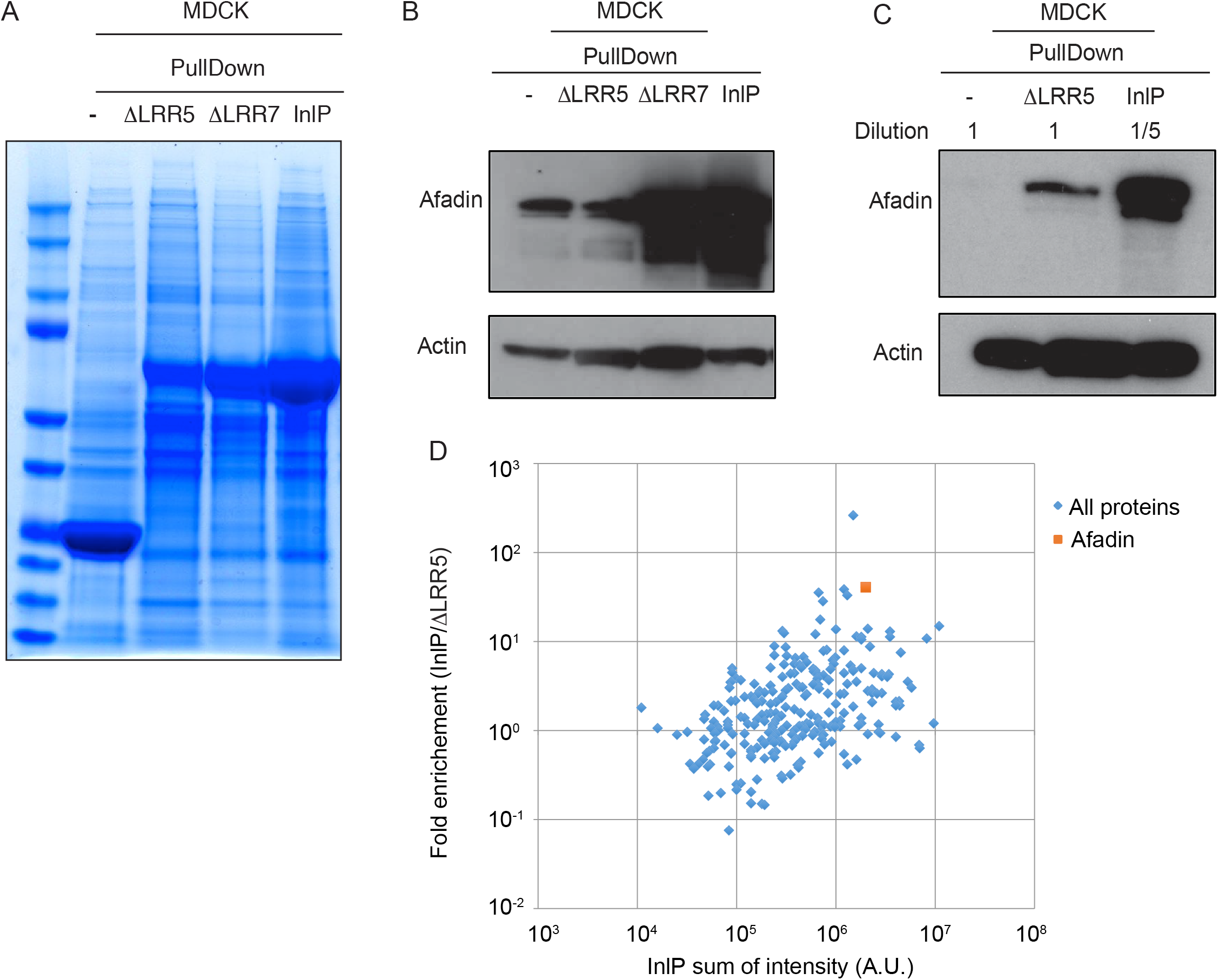
LRR5 stabilized afadin- InlP interaction. (A) Loading control of InlP-afadin binding pull-down experiments with LRR mutants by Coomassie blue staining. GST protein alone (-), InlPΔLRR5-GST fusion protein (ΔLRR5), InlPΔLRR7-GST fusion protein (ΔLRR7), or InlP- GST (InlP) bound to glutathione-Sepharose resin were used as bait for pull-down experiments with protein extracts from MDCK cell line. Data shown are Coomassie staining, and the most abundant band in each lane represents the bait. (B-C) GST fusion proteins or GST protein alone (-) bound to glutathione-sepharose resin were incubated overnight with protein extracts from MDCK cells, and elution fractions (EF) were analyzed by Western blot with anti-afadin antibodies. The following GST fusion proteins were used: GST-InlP (InlP) and GST-InlP with deletions in LRR5 (ΔLRR5) or LRR7 (ΔLRR7). Actin: loading control. Undiluted elution fractions were analyzed with the exception of the elution fractions marked in panel C that were diluted 5-fold. Actin: loading control. (D) Scatter plot of the logarithm of the ratio of intensity of InlP-GST bound proteins to the intensity of ΔLRR5-GST bound proteins versus total intensity of InlP-GST bound proteins. The intensities (A.U.) of the proteins were identified through mass spectrometry using InlP-GST or InlPΔLRR5-GST as baits to identify host binding partners in the MDCK cell culture extracts. Blue diamonds show all the proteins apart from afadin, which is indicated as orange square. Using this metric, afadin is 2.7 standard deviations away from the mean for this group.

To examine the effects of the InlPΔLRR5 mutation on binding to host proteins, we used a GST control, InlP fused to GST (InlP-GST) or InlPΔLRR5 fused to GST (InlPΔLRR5-GST) as baits and identified proteins pulled down from MDCK epithelial cell lysates by mass spectrometry (Fig. 3D and Supplementary Table 2). Of the MDCK cell proteins found specifically to bind InlP- GST, only afadin (*MLLT4/AF-6*) had also been identified in the screens described above using yeast two-hybrid screening on a human placental library (Supplementary Table 1) and mass spectrometry of proteins pulled down from human placental tissue extracts (Supplementary Table 2). Consistent with the Western blot analysis, mass spectrometry analysis confirmed that afadin associated much less strongly with InlPΔLRR5-GST than with InlP-GST (Fig. 3D and Supplementary Fig. 4). These findings are consistent with the hypothesis that the InlPΔLRR5 mutant is specifically deficient in interacting with afadin in MDCK cells, while otherwise retaining much of its normal structure and protein-protein interaction potential.

### Afadin-InlP interactions decrease the attachment of epithelial cells to the extracellular matrix, but not adhesion between adjacent epithelial cells

We sought to investigate whether the interaction between InlP and afadin could alter the structural integrity of epithelial cell monolayers in ways that might facilitate *L. monocytogenes* invasion or cell-to-cell spread. To test whether InlP affects the integrity of cell-cell junctions in a monolayer, we constructed a line of MDCK cells that could express a humanized version of InlP after induction by doxycycline (MDCK-InlP^+^). We cultured wild-type MDCK, MDCK *AF-6*^-/-^, and MDCK-InlP cells on Transwell filters for three days and measured the permeability of the cell monolayers using inulin-FITC. As expected after deletion of a protein necessary for proper formation of adherens junctions and tight junctions (41), MDCK *AF-6*^-/-^ monolayers showed a substantial increase in permeability to inulin relative to wild-type MDCK monolayers. However, MDCK-InlP monolayers showed low permeability, comparable to the baseline level for wild-type MDCK monolayers, both with and without induction of the InlP transgene using doxycycline (Fig. 4A). This result suggests that the interaction between InlP and afadin does not inhibit the ability of afadin to perform its normal function in establishment and maintenance of impermeable cell-cell junctions.

**Figure 4.**
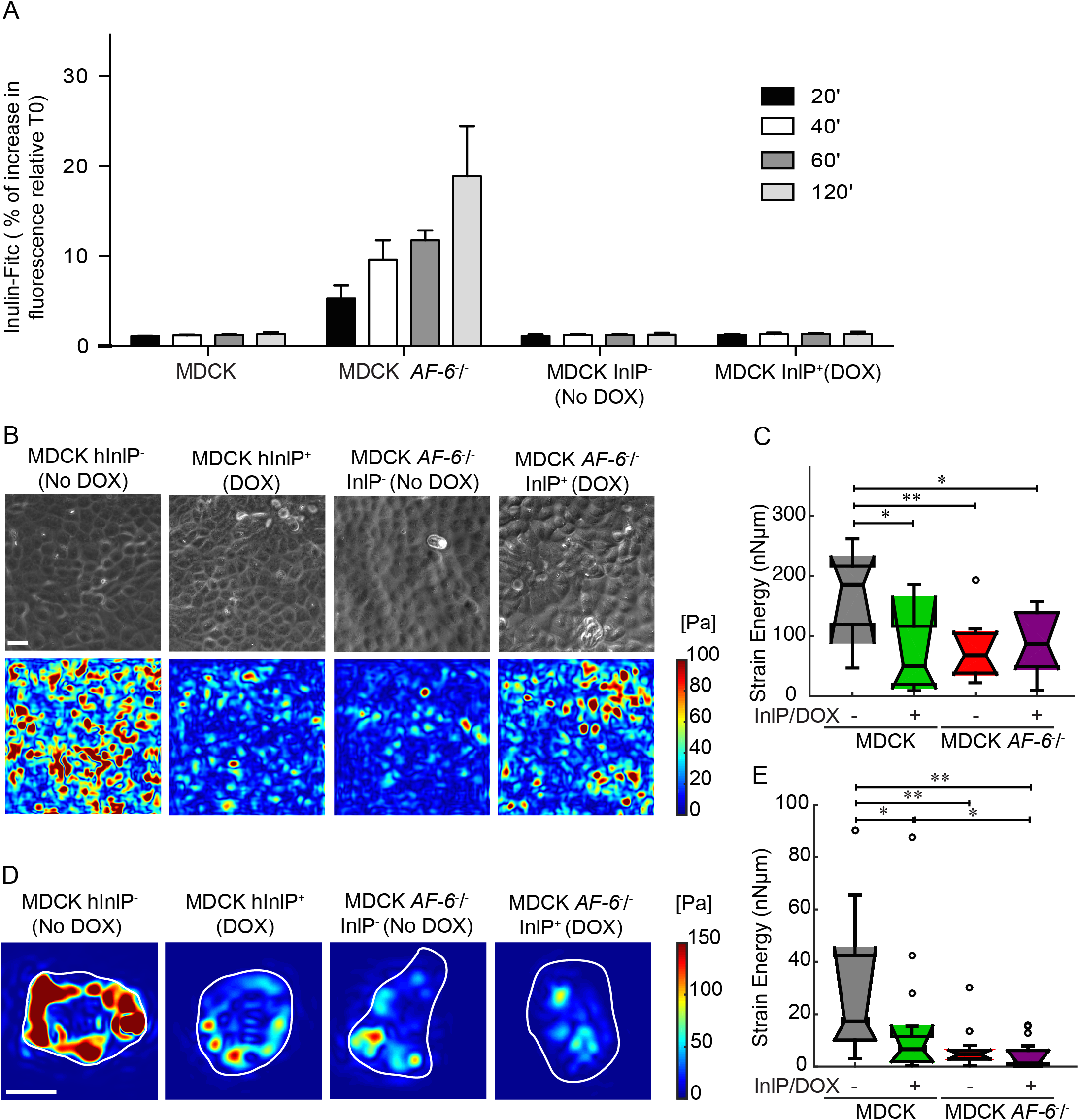
Cell-ECM traction stresses decrease in magnitude when afadin is not expressed or when InlP is present. (A) MDCK, MDCK *AF-6*^-/-^, and MDCK cell lines stably transfected with *inlP* under a doxycycline (DOX) inducible promoter were grown on Transwell filters for three days until formation of polarized epithelia. Inulin-FITC was added to the upper compartment of the Transwell insert. The cell permeability was analyzed at different time points by measuring the fluorescence intensity in the lower compartment. MDCK InlP^-^ and MDCK InlP^+^: stably transfected cells without or with InlP expression secondary to doxycycline exposure for 24 h prior to the addition of Inulin-FITC. (B) Representative phase images (first row) and maps showing the magnitude of traction stresses of confluent MDCK monolayers adherent to hydrogels coated with collagen I (color indicates stress magnitude in Pa). Columns refer to: MDCK InlP^-^, MDCK InlP^+^, MDCK *AF-6*^-/-^ InlP^-^ and MDCK *AF-6*^-/-^ InlP^+^. Scale bar corresponds to 20 *µ*m. (C) Time-averaged strain energy (nN•µm) calculated for multiple regions within confluent MDCK monolayers. Boxplots refer to N=11 MDCK InlP^-^, N=11 MDCK InlP^+^, N=12 MDCK *AF-6*^-/-^ InlP^-^, N= 12 MDCK *AF-6*^-/-^ InlP^+^ monolayer regions. (D-E) TFM results for single cells adherent to polyacrylamide hydrogels coated with collagen I. (D) Instantaneous maps showing the magnitude of traction stresses (color indicates stress values in Pa). Columns refer to: MDCK InlP^-^, MDCK InlP^+^, MDCK *AF-6*^-/-^ InlP^-^ and MDCK *AF-6*^-/-^ InlP^+^. Cell outlines are in white. Scale bar corresponds to 20 *µ*m. (E) Time-averaged strain energy (nNµm) from individual cells. Boxplots refer to N=15 MDCK InlP^-^, N=17 MDCK InlP^+^, N= 15 MDCK *AF-6*^-/-^ InlP^-^, and N=18 MDCK *AF-6*^-/-^ InlP^+^. For boxplots in (C), (E) circles represent outliers, and the boxplots’ notched sections show the 95% confidence interval around the median (Wilcoxon–Mann–Whitney test). One or two asterisks denote statistically significant differences between the medians of two distributions (<0.05 or <0.01, respectively; Wilcoxon rank sum test).

Next, we used these cell lines to examine the modulation of cell-substrate interactions by InlP binding to afadin. To this end, we also generated a line of MDCK *AF-6*^-/-^ cells with the doxycycline-inducible InlP transgene. We used traction force microscopy (TFM) to measure the ability of MDCK cells to generate traction forces on their substrates with and without expression of Afadin and/or InlP. Uninduced MDCK-InlP cell monolayers without doxycycline (the positive control) grown on deformable substrates as confluent, polarized monolayers were able to generate significant strain energy on the order of 100-200 nN·*µ*m (Fig. 4B, C) and to exert traction stresses with peak values around 100 Pa, comparable to previous reports (45, 46). Both the induction of the InlP transgene and the deletion of afadin caused a substantial reduction in strain energy. Induction of InlP expression in the background of the MDCK *AF-6*^-/-^ cell line did not lead to any further reduction, consistent with the hypothesis that the primary effect of InlP expression on abrogation of traction force generation in MDCK cells is due to its interaction with afadin.

We have previously demonstrated that a decrease in monolayer traction force can be caused either by a direct inhibition of cell-substrate interactions or by an indirect inhibition of lateral tension at cell-cell junctions in the monolayer (33). To distinguish between these two possibilities, we also performed TFM for isolated cells under the same four conditions. Just as for the monolayers, either the expression of InlP or the deletion of *AF-6* caused a significant decrease in the ability of isolated individual cells to generate traction force (Fig. 4D, E). Taken together, these results suggest that the InlP-afadin binding interaction specifically disrupts afadin’s normal activity in enhancing traction force generation at the cell-substrate interface but has no effect on afadin’s normal function at cell-cell junctions in the monolayer.

### InlP enhances L. monocytogenes transcytosis through MDCK polarized monolayers

Finally, we examined the influence of the InlP-afadin interaction on infection and spread of *L. monocytogenes* in MDCK monolayers. We infected wild-type and MDCK *AF-6*^-/-^ cells with wild type or *ΔinlP L. monocytogenes*. Gentamicin was added at 30 min to kill extracellular bacteria, and remaining intracellular bacteria were enumerated at 2, 5, 8 and 24 h post-infection. At 2 h, the MDCK *AF-6*^-/-^ cells harbored ~10 times the number of bacteria as compared to wild-type MDCK cells, although the rate of intracellular growth of the bacteria was unchanged. Deletion of *inlP* had no significant effect on invasion or intracellular replication of *L. monocytogenes* in either host cell background (Fig. 5A).

**Figure 5.**
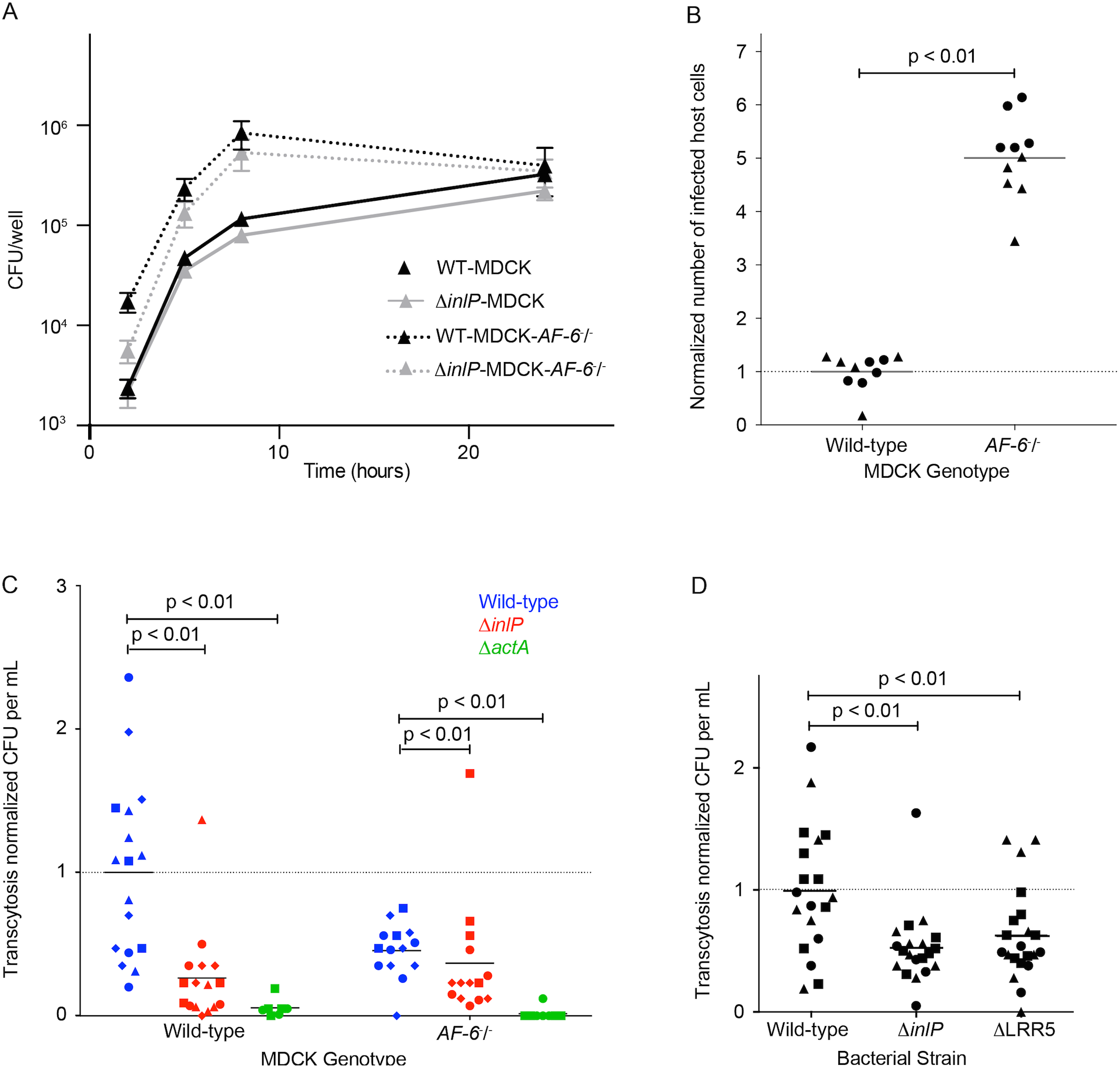
InlP enhances transcytosis through MDCK polarized monolayers. (A) Intracellular growth curves of wild type 10403S *L. monocytogenes* in MDCK (black solid line) or MDCK *AF- 6*^-/-^ (black dashed line) cells and of Δ*inlP L. monocytogenes* in MDCK (gray solid line) or MDCK *AF-6*^-/-^ (gray dashed line) cells. Unpolarized monolayers of MDCK cell lines were infected with 10403S at MOI of 150:1. Gentamicin was added at 50 μg/ml to kill extracellular bacteria and maintained in media thereafter. Each growth curve represents the means and standard deviations of colony forming units (CFU) over time from three separate experiments performed in triplicates. (B) Afadin limits initial *L. monocytogenes* invasion. Flow cytometry quantifying the number of *L. monocytogenes*-containing MDCK and MDCK *AF-6*^-/-^ cells 5 hours post infection. Data were normalized to 1 for wild-type MDCK cells infected with Δ*actA L. monocytogenes* for each experiment, and pooled from two independent experiments (each experiment is depicted by different symbols). (C) Amount of transcytosis by wild-type (blue), Δ*inlP* (red), and Δ*actA* (green) *L. monocytogenes* through MDCK and MDCK *AF-6*^-/-^ monolayers. Data were normalized to 1 for MDCK cells infected with wild-type *L. monocytogenes* for each experiment, and pooled from four independent experiments (MDCK cells) or three independent experiments (MDCK *AF-6*^-/-^ cells). Each experiment is depicted by different symbols. (D) Amount of transcytosis by wild-type, Δ*inlP*, and ΔLRR5 *L. monocytogenes* through MDCK monolayers. Data were normalized to 1 for MDCK cells infected with wild-type *L. monocytogenes* for each experiment, and pooled from three independent experiments. Each experiment is depicted by different symbols.

The observation that MDCK *AF-6*^-/-^ monolayers had substantially more intracellular bacteria at early time points in the course of infection as compared to wild-type MDCK cells raised the possibility that the deletion of *AF-6* caused an increase in the efficiency of *L. monocytogenes* invasion. Because the cell receptor for *L. monocytogenes* invasion into epithelial cells, E-cadherin, is normally sequestered from the apical surface of cells in well-polarized, confluent monolayers, invasion under such circumstances is rare and typically takes place at sites of local, transient disruption of the epithelial cell-cell junctions, for example at sites of extrusion of apoptotic cells (8). Consequently, any disruption of normal cell-cell junctions in epithelial monolayers typically increases the efficiency of *L. monocytogenes* invasion (38). To determine if this was the reason for the increased bacterial load in MDCK *AF-6*^-/-^ monolayers at early time points, we infected both wild-type and *AF-6*-knockout MDCK cells with Δ*actA L. monocytogenes* expressing mTagRFP under the *actA* promoter and counted the number of RFP-positive host cells by flow cytometry 5 h post-inoculation, which represents the number of bacteria-containing cells. Because these bacteria are not capable of actin-based cell-to-cell spread, this assay allows determination of the number of directly invaded host cells, independent of any later variations in bacterial spread (47). Indeed, we found approximately 6-fold more invasion of MDCK *AF-6*^-/-^ cells with *L. monocytogenes* as compared to wild-type MDCKs (Fig. 5B).

Our previous work demonstrated that wild-type and *ΔinlP L. monocytogenes* form foci of similar size in MDCK monolayers (1), suggesting that horizontal cell-to-cell spread in a polarized epithelium is not affected by InlP. This is corroborated by our findings here that the InlP-afadin interaction does not disrupt the function of afadin at cell-cell junctions. However, we also showed that *ΔinlP* strains are impaired in spreading from the CTB epithelial layer into the placental stroma (1), and here we have found that the InlP-afadin interaction specifically disrupts the mechanics of MDCK cell traction force generation at the cell-substrate interface (Fig. 4). We therefore wondered whether this protein-protein interaction might specifically affect the ability of intracellular *L. monocytogenes* to use actin-based motility to spread through the basal face of the CTB epithelial monolayer underlying the placental STB, rather than horizontally through the lateral faces of neighboring CTB epithelial cells. We designed an assay to mimic this particular type of spread by growing MDCK cells on Transwell filters with large (3 *µ*m) pores through which bacterial protrusions could extend into the lower well. Under these culture conditions, MDCK cells form well-polarized monolayers and secrete a thick and well-organized basement membrane (basal lamina) (48), so bacterial protrusions reaching the lower well must have been able to form at the basal side of the cell and push through the basement membrane. Approximately 5 hours after inoculation from the apical side, bacteria that had successfully performed transcytosis through the monolayer were collected from the lower well. In this assay, *ΔactA* bacteria exhibited a substantial (>10-fold) reduction in transcytosis, demonstrating that this process is dependent on bacterial actin-based motility (Fig. 5C). The *ΔinlP* strain also showed a statistically significant reduction in transcytosis (~4-fold), consistent with the hypothesis that the InlP-afadin interaction specifically alters cell mechanics at the basal cell-substrate interface and thereby facilitates *L. monocytogenes* spread in a direction perpendicular to the monolayer.

If the nature of the InlP-afadin interaction were simply to sequester or inhibit afadin’s normal activity at the basal surface, we would expect that deletion of *AF-6* in the host cells should be able to rescue the transcytosis defect of *ΔinlP L. monocytogenes*. Interestingly, however, this was not the case. MDCK *AF-6*^-/-^ monolayers infected with wild-type bacteria exhibited an unexpected decrease in transcytosis efficiency, resulting in about two-fold fewer bacteria reaching the lower well as compared to parallel infections in wild-type MDCK cells, despite the much higher efficiency of initial invasion (Fig. 5C). In this host cell background, transcytosis of *ΔinlP* bacteria was slightly less efficient than that of wild-type bacteria, in contrast to the expected phenotypic rescue.

We wished to confirm that this unexpected observation could be attributed specifically to the InlP-afadin interaction, rather than to some pleiotropic effect of *AF-6* deletion in the MDCK cells, and so took advantage of our earlier determination that the InlPΔLRR5 mutation specifically disrupts afadin binding (Fig. 3 and Supplementary Table 2). Infection of wild-type MDCK cells with *L. monocytogenes* expressing the mutant InlPΔLRR5 resulted in a transcytosis defect comparable to the defect of the complete *ΔinlP* deletion in parallel experiments (Fig. 5D). We conclude that the InlP-afadin protein-protein interaction exerts a specific effect to enhance transcytosis of *L. monocytogenes* through the basal face of polarized epithelial cells that is not equivalent to the simple removal or sequestration of afadin from this site.

## DISCUSSION

En route to infection of the fetus, *L. monocytogenes* experiences multiple bottlenecks, from trafficking to the placenta (3) to surviving the placental innate immune defenses (49) to crossing the trophoblast monolayer and its associated basement membrane into the fetal stroma and fetal circulation (20). InlP initially attracted our interest due to its identification in a screen for *L. monocytogenes* mutants that were defective in their ability to infect the placenta (1). Two unbiased screening methods to identify potential placental binding partners, a yeast two-hybrid screen using a placental cDNA library and mass spectrometric identification of human proteins pulled down by InlP from placental extracts, converged on afadin as a significant candidate binding partner. Afadin is not found on the cell surface, but instead is primarily associated with the cytoplasmic face of cell-cell junctions containing nectin (41, 53). Therefore, it seems likely that InlP is most helpful in breaching that third barrier and accessing the fetal stroma.

Of the 25 known members of the *L. monocytogenes* internalin family, the only other one known to have host cell cytoplasmic binding partners involved in its virulence functions is InlC. InlC binds to one of the subunits of the IkB complex thereby modulating the host cell innate immune response (54). In addition, InlC also binds the adapter protein Tuba, which cooperates with the actin polymerization regulator N-WASP to modulate tension at cell-cell junctions and thereby promote *L. monocytogenes* cell-to-cell spread in epithelial monolayers (55). Intriguingly, InlC and InlP are two of only four internalin family members that are thought to be secreted in a soluble form rather than anchored to the bacteria cell surface (24). As more internalin family members are characterized, it will be interesting to see whether the trend persists of surface-anchored internalins binding to host cell surface proteins and secreted internalins binding to host cell cytoplasmic proteins.

While we have not determined the structure of the InlP-afadin complex, our work does give some clues into the likely nature of this interaction. The internal in-frame deletion mutant InlP-ΔLRR5, which lacks one of the leucine-rich repeats, binds afadin less well than full length InlP, but this is probably not simply due to the shorter overall size of the concave face of the LRR domain because InlP-ΔLRR7 and InlP-ΔLRR8, which should have a similar overall length of the LRR domain concave face, seem to have little or no deficiency in afadin binding. It is thus likely that there are specific residues in LRR5 that stabilize the interaction. An interesting unanswered question concerns the role of the unusual protruding loop from LRR3 in determining binding partner specificity. Although we found that this loop between amino acids 132-136 in the InlP structure bound Ca^2+^ with an affinity of 35 μM, this binding affinity is probably too weak to be relevant in the host cell cytoplasm where the Ca^2+^ concentration rarely rises above ~2 μM (56). This is not the case for other subcellular compartments like the acidifying phagosomes, where the Ca^2+^ concentration has been shown to be >70 μM (56). Thus, it is possible that calcium binding by InlP plays a role during *L. monocytogenes* escape from the primary (single-membrane) or secondary (double-membrane) vacuole. It should be noted that Shaughnessy et al. (57) observed that listeriolysin O pores generated by the bacterium in the process of primary vacuolar escape allow Ca^2+^ in the vacuole to leak into the cytoplasm, albeit slowly. It is possible that calcium binding promotes changes in InlP behavior that are specific to different calcium microdomains in the polarized epithelium (58, 59), including microdomains potentially specific to basal protrusions and/or placental trophoblasts (60).

Other bacterial pathogens are also known to secrete virulence factors into the host cell cytoplasm that modulate host cell biology in ways that benefit bacterial survival or spread (33, 61-64). One particularly relevant example is the virulence factor Sca4, secreted by *Rickettsia parkeri*. Analogous to InlC, Sca4 binds the host protein vinculin at cell-cell junctions and alters cortical tension, promoting actin-based cell-to-cell spread of the bacterium horizontally through host cell monolayers (33). Given the precedent that both InlC and Sca4 bind to proteins that perform scaffolding functions at cell-cell junctions in polarized epithelia (Tuba and vinculin respectively) and modulate cortical tension to promote bacterial spread, our initial expectation on discovering that InlP binds the cell-cell junction-associated protein afadin was that its mechanism of action would be similar. However, we have found no evidence that InlP has any effect on cell-cell junctions in the host monolayer, or indeed that it makes any significant contribution to lateral actin-based spread of *L. monocytogenes* within the horizontal plane of the monolayer.

Instead, our results indicate that the relevant site of action of InlP in the host cell is the basal face, not the lateral face, and that the InlP-afadin interaction specifically modulates the nature of the host cell organization at the basal face or interaction with the underlying basement membrane in a way that promotes *L. monocytogenes* spread that is perpendicular, rather than parallel to the monolayer. Afadin has not been described as being localized to the basal face of polarized epithelial cells, or to cell-ECM junctions; instead its functional characterization has focused primarily on nectin binding at cell-cell junctions (40, 41). Conditional knockouts in the mouse intestine lead to increases in intestinal permeability (65), consistent with the phenotype we have described here of afadin loss causing permeabilization of cell-cell junctions in MDCK cells. Whether afadin also plays a role in cell-ECM connections is unclear, but in certain cell types, removal of afadin can enhance cell migration (66) and invasive capacity (67), or suppress neuronal axon branching (68). While these cellular behaviors might involve specific changes in cell-ECM interactions, it is also likely in these cases that the effects of afadin disruption are mediated indirectly through its interactions with critical signaling molecules, Rap1, Src and R-Ras respectively (66-68).

How does InlP binding modulate afadin activity at the basal face of epithelial cells? For both InlC and Sca4, deletion of the host cell binding partner is sufficient to (partially) rescue the bacterial deletion phenotype, consistent with the simple idea that binding of the bacterial virulence factors to their host cell partners effectively inhibits or sequesters those host cell partners (33, 55). In contrast, our results indicate that the consequences of the interaction between InlP and afadin are more complex than simple sequestration. InlP expression in uninfected MDCK cell monolayers has no effect on integrity of cell-cell junctions, while deletion of afadin results in a significant increase in monolayer permeability, so the presence of InlP does not appear to disrupt afadin’s normal activities at the cell-cell junctions. In contrast, expression of InlP phenocopies afadin deletion with respect to the ability of epithelial cells to generate traction forces on their substrates. And most intriguingly we have found that *AF-6* knockout MDCK cells are less able to support transcytosis of both wild-type and *ΔinlP L. monocytogenes* across the basement membrane than are wild-type host cells; if afadin normally simply strengthens the cell-substrate interface then the *AF-6* knockout host cells should be more permissive for bacterial transcytosis, rather than less. This raises the interesting possibility that the InlP-afadin binding interaction results in a novel activity that specifically promotes bacterial formation of actin-based protrusions at the cell basal face.

In the context of placental infection, the spatial specificity for the activity of the InlP- afadin interaction at the basal face of polarized epithelial cells is relevant, as our previous work has suggested that vertical spread from the CTB monolayer into the underlying fetal stroma is a major barrier for infection of the fetus (1, 20, 69). However, we previously showed that the Δ*inlP* mutant is not impaired in reaching the liver or spleen in orally-infected guinea pigs (1), raising the question of why the phenotype is apparently specific for placental infection when enhanced transcytosis through the basement membrane should also be relevant for *L. monocytogenes* crossing the intestinal epithelium or crossing the endothelial cells in blood vessels to invade the liver. In this context, we note that the intestinal barrier is organized differently from that of the placenta. The intestinal epithelium is rich in intercellular junctions that link neighboring cells together, while the placental syncytiotrophoblast that contacts maternal blood is a single, continuous plasma membrane extending over tens of thousands of square millimeters in later stages of human gestation. The invasive cytotrophoblasts in the uterine decidua do have exposed lateral surfaces and cell-cell junctions, but they lack the lymphatic and immune components of the intestine in that there are no Peyer’s Patches and M cells with phagocytic and migratory capabilities. While some white blood cells do traffic across this barrier in both directions (70), traffic is likely more limited than in the mucosal-associated lymphoid tissues of the gut. The lack of a role for InlP in crossing the intestinal barrier is therefore consistent with earlier work showing that *L. monocytogenes* traffics from the gut to the bloodstream by being carried by immune cells (71). In this context, the limited immune cell traffic observed across the placental barrier might explain how Δ*inlP L. monocytogenes* are impaired in crossing the subsyncytial CTB monolayer— undergirded by a basement membrane—into the fetal stroma (1). Alternatively, it is possible that the basement membrane underlying the CTB layer simply presents a more formidable physical barrier than that found underneath the intestinal epithelium. An interesting issue, to be addressed in future work, is whether there is impairment in the invasion of *ΔinlP L. monocytogenes* across the blood-brain endothelial barrier.

For decades, *L. monocytogenes* has been a useful tool for the study of mammalian cells, barriers, and immune functions. Increasingly, its capacity to reveal basic knowledge about higher levels of biological organization—mammalian tissues and organs—is becoming evident. Through investigations of how bacteria cross barriers and of the interactions between bacterial factors and host pathways, we learn not only more about what regulates pathogenesis, but about the structures of these tissues themselves, in how they are generated, their composition, and how they can be disrupted.

## AUTHOR CONTRIBUTIONS

C.F. and A.I.B. conceived the project and initiated collaborations with the other authors. C.F. generated all bacterial mutant strains and mammalian cell lines. C.F., E.E.B., F.E.O., S.H.L., G.R., and S.N. performed experiments and/or analyzed data. W.F.A., J.A.T. and A.I.B. provided resources and supervision. C.F., E.E.B., J.R.R., A.I.B. and J.A.T. wrote the paper.

## ACKNOWLEDGEMENTS

Our thanks to Henriette Macmillan, Cedric Brimacombe, George Minasov, Olga Kiryukhina and Albin Cardona-Correa for discussions and experimental support, as well as tissue donors who provided informed consent. This work was supported by NIH R01AI084928 (A.I.B), Burroughs Wellcome Fund (A.I.B), NIH R01AI036929 (J.A.T), HHMI (J.A.T), the HHMI Gilliam Fellowship for Advanced Study (F.E.O.), the Stanford Graduate Fellowship (F.E.O.), the American Heart Association (E.E.B.), NIH F32AI108195 (G.R.), Society for Pediatric Pathology Young Investigator Research grant (G.R.), and the University of California Partnerships for Faculty Diversity President’s Postdoctoral Fellowship (G.R.). Flow cytometry was performed at the Stanford Shared FACS Facility. The Center for Structural Genomics of Infectious Diseases has been funded with federal funds from the National Institute of Allergy and Infectious Diseases, National Institutes of Health (NIH), Department of Health and Human Services, under Contract Nos. HHSN272200700058C and HHSN272201200026C (to W.F.A). This research used resources of the Advanced Photon Source, a U.S. Department of Energy (DOE) Office of Science User Facility operated for the DOE Office of Science by Argonne National Laboratory under Contract No. DE-AC02-06CH11357. Use of the LS-CAT Sector 21 was supported by the Michigan Economic Development Corporation and the Michigan Technology Tri-Corridor (Grant 085P1000817).

## SUPPORTING INFORMATION CAPTIONS

**Figure S1. Coomassie and silver stains showing fractions for mass spectrometry analysis**. GST protein alone and InlP -GST fusion protein bound to glutathione-Sepharose resin were used as bait for pull-down experiments with (+) or without (-) protein extracts from human placenta. Shown on the left is Coomassie blue staining of each fraction, with the most abundant band in each lane representing the bait protein. Shown on the right is silver staining of each fraction, with the most abundant band in each lane representing the bait protein. The red boxes labeled A and B show the two fractions extracted from gel and analyzed by mass spectrometry (see Sheet 1 and 2 in Supplementary Table 2).

**Figure S2. Structural and biochemical details on InlP Ca^2+^ interaction**. (A) Schematic view of the interaction between the Ca^2+^ with the 3^rd^ LRR loop of InlP. The amino acids D132 and E182, which are involved in the interaction, are shown as sticks, water molecules as red spheres, and hydrogen bonds as dashed lines. (B) Isothermal titration calorimetry results show Ca^2+^ binding to InlP. The isotherm was fit by a one site binding model (N = 1.8 ± 0.1 site, K = 2.6E4 ± 6.5E3 M^-1^, ΔH = -1176 ± 90.2 cal/mol, ΔS = 16.3 cal/mol/deg).

**Figure S3. Loading control by Coomassie staining and western blot analysis of pull down experiments on InlPΔLRR8 mutant**. InlP-afadin binding pull-down experiments with ΔLRR8 mutant. GST protein alone (-), InlPΔLRR8-GST fusion protein (ΔLRR8), or InlP-GST (InlP) bound to glutathione-sepharose resin were used as bait for pull-down experiments with protein extracts from MDCK cell line. (A) Coomassie blue staining of each fraction, with the most abundant band in each lane representing the bait protein. (B) Data shown are a western blot analysis using anti-afadin antibodies. Actin was used as loading control.

**Figure S4. LRR5 stabilized afadin- InlP interaction**. Scatter plot of intensities of InlP-GST versus ΔLRR5-GST binding proteins coming from MDCK cell cultures extracts. Plot shows the sum of intensities (A.U.) of the proteins identified through mass spectrometry using InlP-GST or InlPΔLRR5-GST as baits to identify host binding partners in the MDCK extracts. Blue diamonds show all the proteins apart from afadin, which is indicated as orange square. Black line shows X=Y. Filters applied to the data are discussed in the Methods section.

## SUPPLEMENTARY TABLES

**Supplementary Table 1. Yeast two-hybrid screening of human placenta library**.

**Supplementary Table 2. Mass spectrometry data from pull-down of InlP-GST interacting with binding partners from human placental tissue and from extracts from MDCK cell cultures**.

**Supplementary Table 3. Crystallographic Table**.

---

**Table 1.** Yeast two-hybrid screening of human placenta library. Results summary listing the protein partners identified (gene name), the number of independent clones and the global PBS ^a^. Ordering is performed based on global PBS and then on number of clones. Afadin is indicated in red.

| Gene name | Number of independent clones | Global PBS<sup>a</sup> |
| --- | --- | --- |
| RPS8 | 26 | A |
| RBM5 | 14 | A |
| TSHZ1 | 12 | A |
| RPL5 | 9 | A |
| AEBP1 | 7 | A |
| NAB2 | 6 | A |
| THAP3 | 6 | A |
| NAB1 | 4 | A |
| DNMT3A | 5 | B |
| SAFB | 5 | B |
| ZNF653 | 5 | B |
| EXOSC3 | 4 | B |
| RPL7A | 4 | B |
| AF6 | 3 | B |
| FLJ22329 | 3 | B |
| SP3 | 3 | B |
| ZNF653 | 5 | B |
| ZNF6 | 3 | C |
| CDC27 | 2 | C |
| EPS8L2 | 2 | C |
| PHF23 | 2 | C |
| ZNF283 | 2 | C |

<sup>a</sup>Global Predicted Biological Score (PBS). Confidence score of each interaction. A: very high confidence interaction; B: high confidence in the interaction; and C: good confidence in the interaction.

**Table 2.**
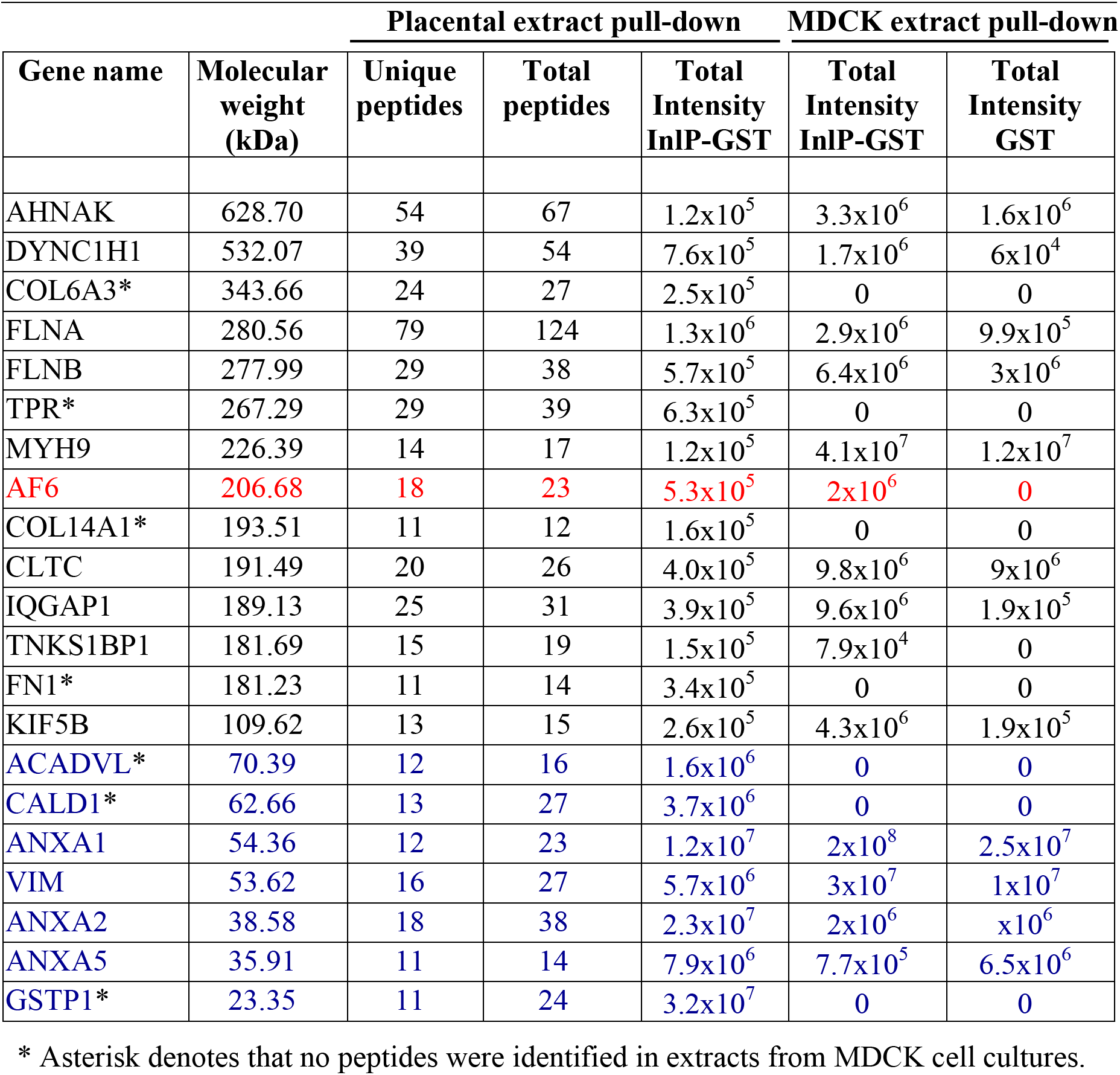
Mass spectrometry data from pull-downs of InlP-GST. Proteins identified with more than 10 unique peptides from human placental extract pull-downs are outlined below and ordered based on descending molecular weight. Black and blue gels correspond to sections A and B respectively (see Supplementary Figure 1). Afadin is indicated in red. Columns 3-5 correspond to pull-downs from placental extract with InlP-GST. Columns 6-7 show the corresponding total intensities from mass spectrometry data using InlP-GST and GST alone as baits to identify host binding partners in extracts from MDCK cell cultures.

**Table 3.** Crystallographic Table. Data Collection and Refinement Statistics^a^

|  | **InlP** | **Lmo2027** |
| --- | --- | --- |
| *PDB code* | 5HL3 | 5KZS |
| *Data Collection* |  |  |
| Resolution range, Å | 30.00-1.40<br>(1.42-1.40) | 30.00-2.30<br>(2.34-2.30) |
| Space group | P4<sub>1</sub>2<sub>1</sub>2 | I<sub>4</sub> |
| Unit cell dimensions |  |  |
| a,b,c, Å | a & b = 72.9<br>c = 179.5 | a & b = 157.8<br>c = 36.3 |
| α, β, γ, degrees | α, β, γ = 90 | α, β, γ = 90 |
| Completeness, % | 99.8 (100) | 98.6 (99.6) |
| No. reflections | 180423 (9031) | 20005 (1429) |
| Redundancy | 45.6 (5.3) | 6.2 (4.5) |
| ⟨I/σ(I)⟩ | 30.2 (2.6) | 79.7 (2.0) |
| R<sub>merge</sub>, % | 7.9 (65.3) | 9.0 (57.1) |
| *Refinement statistics* |  |  |
| R-factor (R<sub>work</sub>/R<sub>free</sub>)<sup>b</sup> | 16.0/17.6 | 22.8/27.9 |
| No. atoms |  |  |
| Protein | 2935 | 2377 |
| Waters | 828 | 26 |
| Ca<sup>2+</sup> | 8 | n/a |
| R.M.S. deviations |  |  |
| Bond lengths, Å | 0.011 | 0.009 |
| Bond angles, degrees | 1.51 | 1.47 |
| Ramachandran analysis |  |  |
| Favored regions, % | 96 | 95 |
| Allowed regions, % | 100 | 99 |
| Disallowed regions, % | 0 | 1 |

<sup>a</sup> Highest resolution shell in parenthesis.<sup>b</sup> Definition of R<sub>work</sub>, R<sub>free</sub>: $R = \sum_{hkl} ||F_{\text{obs}}| - |F_{\text{calc}}|| / \sum_{hkl} |F_{\text{obs}}|$ , where *hkl* are the reflection indices used in refinement for **R<sub>work</sub>**, and **5% not used in refinement for R<sub>free</sub>**. *F<sub>obs</sub>* and *F<sub>calc</sub>* are structure factors deduced from measured intensities or calculated from the model, respectively.

